# Weighted minimizer sampling improves long read mapping

**DOI:** 10.1101/2020.02.11.943241

**Authors:** Chirag Jain, Arang Rhie, Haowen Zhang, Claudia Chu, Sergey Koren, Adam Phillippy

**Affiliations:** National Human Genome Research Institute, National Institutes of Health, Bethesda, MD 20892, USA; College of Computing, Georgia Institute of Technology, Atlanta, GA 30332, USA

## Abstract

**Motivation:** In this era of exponential data growth, minimizer sampling has become a standard algorithmic technique for rapid genome sequence comparison. This technique yields a sub-linear representation of sequences, enabling their comparison in reduced space and time. A key property of the minimizer technique is that if two sequences share a substring of a specified length, then they can be guaranteed to have a matching minimizer. However, because the *k*-mer distribution in eukaryotic genomes is highly uneven, minimizer-based tools (e.g., Minimap2, Mashmap) opt to discard the most frequently occurring minimizers from the genome in order to avoid excessive false positives. By doing so, the underlying guarantee is lost and accuracy is reduced in repetitive genomic regions.

**Results:** We introduce a novel weighted-minimizer sampling algorithm. A unique feature of the proposed algorithm is that it performs minimizer sampling while taking into account a weight for each *k*-mer; i.e, the higher the weight of a *k*-mer, the more likely it is to be selected. By down-weighting frequently occurring *k*-mers, we are able to meet both objectives: (i) avoid excessive false-positive matches, and (ii) maintain the minimizer match guarantee. We tested our algorithm, Winnowmap, using both simulated and real long-read data and compared it to a state-of-the-art long read mapper, Minimap2. Our results demonstrate a reduction in the mapping error-rate from 0.14% to 0.06% in the recently finished human X chromosome (154.3 Mbp), and from 3.6% to 0% within the highly repetitive X centromere (3.1 Mbp). Winnowmap improves mapping accuracy within repeats and achieves these results with sparser sampling, leading to better index compression and competitive runtimes.

**Availability:** Winnowmap is built on top of the Minimap2 codebase (Li, 2018) and is available at https://github.com/marbl/winnowmap.

## 1 Introduction

Continued development of time and space-efficient algorithmic techniques has been pivotal for dealing with the exponential growth of DNA sequencing throughput. In the context of mapping and alignment applications, tools have evolved from purely alignment-based (Smith and Waterman, 1981), to seed-and-extend (Altschul *et al.*, 1997; Kurtz *et al.*, 2004), to succinct text indexing (Langmead and Salzberg, 2012; Yu *et al.*, 2015), and now to ‘sketch’-based (Ondov *et al.*, 2016; Li, 2018) techniques. Sketch-based algorithms employ dimensionality reduction to transform a sequence into a more compact representation, e.g., a subset of *k*-mers present in the sequence (Marçais *et al.*, 2019; Rowe, 2019). While these algorithms continue to be widely leveraged in bioinformatics, they are even more prevalent for long-read (PacBio/Oxford Nanopore) analyses because longer strings are more amenable to compaction. As such, several long-read based mappers (Li, 2016; Popic and Batzoglou, 2017; Jain *et al.*, 2018; Li, 2018), genome assemblers (Berlin *et al.*, 2015; Koren *et al.*, 2017; Chin and Khalak, 2019; Shafin *et al.*, 2019; Kundu *et al.*, 2019), metagenomic read classifiers (Dilthey *et al.*, 2019), transcriptomic tools (Sahlin and Medvedev, 2019; Sahlin *et al.*, 2020) employ either minimizer- or MinHash-based sequence comparison.

Minimizer sampling was introduced to the field by Roberts *et al.* (2004) for scaling the genome assembly problem, after being independently described a year earlier by Schleimer *et al.* (2003) in the text mining literature. Given a fixed window length *w* and a pre-defined ordering of all *k*-mers, minimizer sampling selects the minimum *k*-mer from every consecutive window (Figure 1). Minimizers can be collected in linear time with regard to the sequence length, and matching substrings can be quickly identified since the algorithm is guaranteed to select a common minimizer for two sequences if they share an exact match of at least *w* + *k* − 1 bases long. Further, if the ordering of *k*-mers is determined using a random permutation, then the expected minimizer sampling density is known to be 2*/*(*w* + 1) (Roberts *et al.*, 2004). Due to these properties, there exists a trade-off between sensitivity and speed when deciding the window length *w*.

**Fig. 1.**
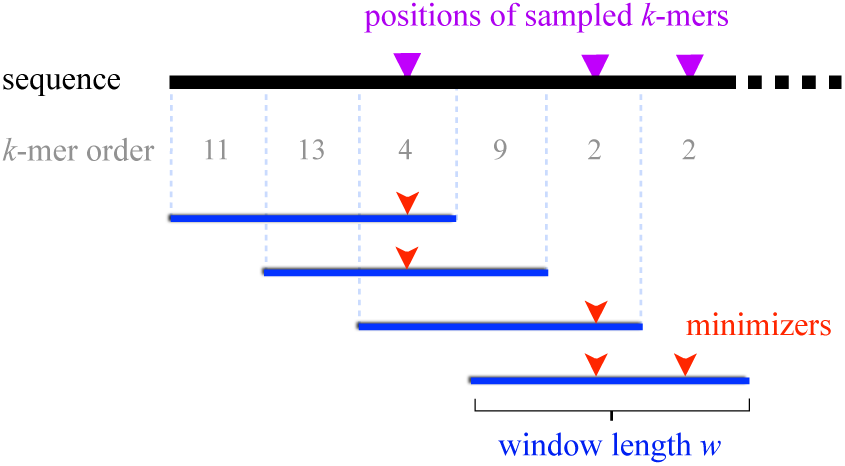
Visualization of the minimizer sampling technique (Roberts et al., 2004). The sequence being sampled is shown as a black line. Assuming a pre-defined order of *k*-mers (e.g., lexicographical), the sampling selects each of the smallest *k*-mer(s) (shown as red arrows) in consecutive sliding windows (shown as blue intervals). The length of each window, i.e., *w* is three in the above example.

As one would expect, minimizers originating from repetitive genomic segments are sampled at a higher frequency than minimizers from unique regions. When used for seeding pairwise alignments, high-frequency minimizers lead to excessive false-positives and can prohibitively increase runtime and memory usage. A popular yet sub-optimal way to deal with this problem is to mask minimizers whose frequency exceeds a certain threshold. For instance, Minimap2 (Li, 2018) discards the top 0.02% most frequently occurring minimizers, whereas Mashmap (Jain *et al.*, 2018) discards the top 0.001% by default. An unfortunate consequence of masking is that minimizer matches are no longer guaranteed and accuracy is reduced in repetitive genomic regions. Negative effects of *k*-mer masking have been highlighted before, such as poor homology detection in genomic repeats (Frith, 2011) and mis-assembly of high-copy plasmids (Koren *et al.*, 2017).

To address the above problem, we propose optimizations to the standard minimizer sampling technique. We formulate the idea of ‘weighted-minimizers’, where *k*-mers that are assigned higher (or lower) weights are more (or less) likely to be selected as minimizers. Using this technique, we show that down-weighting highly repetitive *k*-mers avoids excessive false-positive matches, while still maintaining the guarantee that all matching substrings of length *w*+*k*−1 or greater will share a minimizer. In addition to weighted minimizers, we also incorporate and demonstrate the utility of ‘robust winnowing’ (Schleimer *et al.*, 2003). In the case of multiple equally-minimum *k*-mers in a window, robust winnowing breaks the tie by preferring a *k*-mer that has already been chosen by a previous window. This differs from the standard algorithm (Roberts *et al.*, 2004), which selects all equally-minimum *k*-mers, and is a simple yet crucial trick for efficient handling of low-complexity sequences such as “ACACAC…”. After implementing the above optimizations, we prove that the time complexity to compute weighted, robust minimizers remains linear with regard to sequence length and that the original minimizer match guarantee is preserved.

We refer to our implementation as Winnowmap, which replaces the minimizer sampling and indexing algorithms of the widely used alignment tool Minimap2 (Li, 2018). In doing so, we reuse Minimap2’s highly efficient anchor chaining and gapped alignment routines. We evaluate the speed and accuracy of Winnowmap using simulated and real long-read sequencing data aligned to the human reference genome GRCh38 (Schneider *et al.*, 2017), including the recently completed X chromosome that contains the first-ever assembled human centromere (Miga *et al.*, 2019). Compared to Minimap2, our results demonstrate that Winnowmap reduces the mapping error-rate over the entire X chromosome (0.14% to 0.06%), with the biggest gains achieved in the highly repetitive centromeric region (3.6% to 0.0%). Maintaining high alignment accuracy within long genomic repeats is critical for accurate genome assembly and variant calling. By avoiding masking, we show that Winnowmap maintains uniform minimizer density, while the masking heuristic used by Minimap2 leads to significant drops in minimizer density, especially within long tandem repeats like the centromeric satellite array. Moreover, Winnowmap uses less memory (up to 50% less) while maintaining a similar runtime versus Minimap2. Here we focus on sequence read mapping, but we expect our optimizations to benefit all applications of minimizers.

### Related work

Once a *k*-mer ordering is defined, the set of minimizers picked from an input sequence becomes deterministic. Accordingly, our idea of weighted-minimizers is implemented by manipulating the *k*-mer ordering to prefer higher weighted *k*-mers. There have been prior attempts to optimize minimizer selection for different applications. Chikhi *et al.* (2015) use minimizers for low-memory construction and traversal of de Bruijn graphs. In their algorithm, a complete set of *σ*^*k*^ *k*-mers (*σ* is the size of the alphabet) is sorted based on *k*-mer frequency to define an order. Accordingly, a low-frequency *k*-mer is preferred over a high-frequency *k*-mer in each window. This procedure, however, requires 64 × *σ*^*k*^ bits in memory. This would work for small values of *k* (≤ 10 in their application), but does not scale to the larger *k*-mer sizes (e.g. 15–21) typically used for long-read analyses. Other work has focused on optimizing *k*-mer ordering to reduce the minimizer density (Orenstein *et al.*, 2016; Marçais *et al.*, 2018; DeBlasio *et al.*, 2019). These optimizations are complementary to our objective as we seek to avoid excessive false-positive hits that occur by sampling high frequency *k*-mers. Our formulation of weighted-minimizers is partly inspired by (Chum *et al.*, 2008), which extends the MinHash technique to incorporate *tf-idf* weighting and was adopted by Canu (Koren *et al.*, 2017) for sequence read overlap detection. Below we present background on the classic minimizer sampling (Roberts *et al.*, 2004) and weighted MinHash (Chum *et al.*, 2008) techniques, the two key ideas upon which our algorithm is based.

## 2 Background

### 2.1 Minimizer sampling

Here we formally discuss the standard minimizer sampling algorithm and recall its properties. Continuing the same notation, *w* denotes window length and *σ* is size of the alphabet. Let *U* be the universe of all *σ*^*k*^ *k*-mers, and *h* : *U* → [0, 1] be a random hash function that assigns each *k*-mer to a real number within a unit interval. The function *h* induces an ordering among *k*-mers. Although Roberts *et al.* (2004) originally described the algorithm using a lexicographical ordering, a randomized ordering often works better in practice (Marçais *et al.*, 2017).

The minimizer sampling algorithm entails computing the minimum *k*-mer within each consecutive window of length *w*. Ties are handled by picking all equally minimum *k*-mers (Figure 1). Perhaps the easiest way to compute minimizers is to use two loops such that the outer loop iterates through the input sequence while the inner loop computes a minimum *k*-mer within each window. Assuming the hash *h* of a *k*-mer is computable in *O*(1) time, this nested-loop approach requires *O*(*nw*) time for an input sequence of length *n*. However, there exists a linear-time *O*(*n*) algorithm for computing sliding-window minimum elements (Smith, 2011), where the trick is to use a double-ended queue while streaming *k*-mers (Figure 2). This algorithm starts with an empty queue *Q* (Algorithm 1, line 1). Before inserting a new *k*-mer at the back end of *Q*, the algorithm scans and discards all *k*-mers higher than the current one (line 5). This is because all the previously seen higher *k*-mers cannot be minimizers of either the current or subsequent windows. Due to this step, *Q* maintains sorted *k*-mer order in each iteration. After inserting the new *k*-mer, we also discard those *k*-mers from the front that are located outside the range of the current window (line 8). Due to the sorted order in *Q*, the smallest *k*-mer is simply drawn from front of *Q* (line 12). Note that each element of the array *arr* is inserted and deleted only once throughout the algorithm. Therefore, the total runtime of the algorithm is linear in sequence length. Schleimer *et al.* (2003) proved that the expected density, i.e. fraction of *k*-mers sampled from a random input sequence equals 2*/*(*w* + 1) for all practical values of *w* (*w* < *σ*^*k*^).

**Fig. 2.**
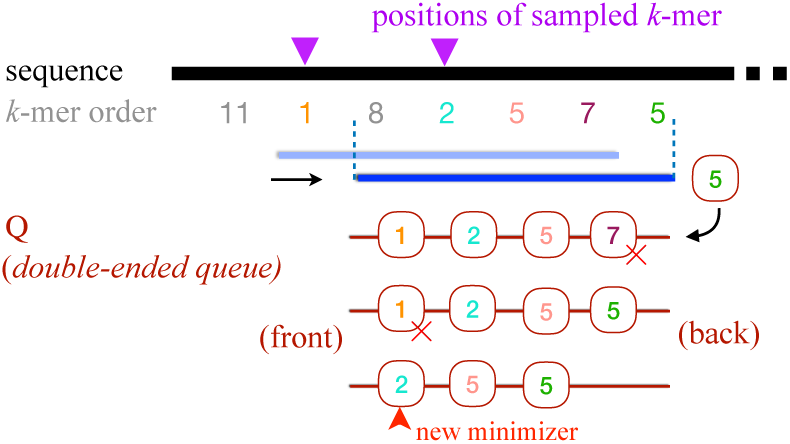
Visualizing a single iteration of Algorithm 1. As the window slides to the right, the new *k*-mer is inserted at the back of the double-ended queue *Q* and a minimizer is selected from front end of the queue. The above figure also uses two equal *k*-mers (order= 5) in different colors to highlight how ties are handled in Algorithm 1.

#### Algorithm 1: Standard procedure for computing minimizers

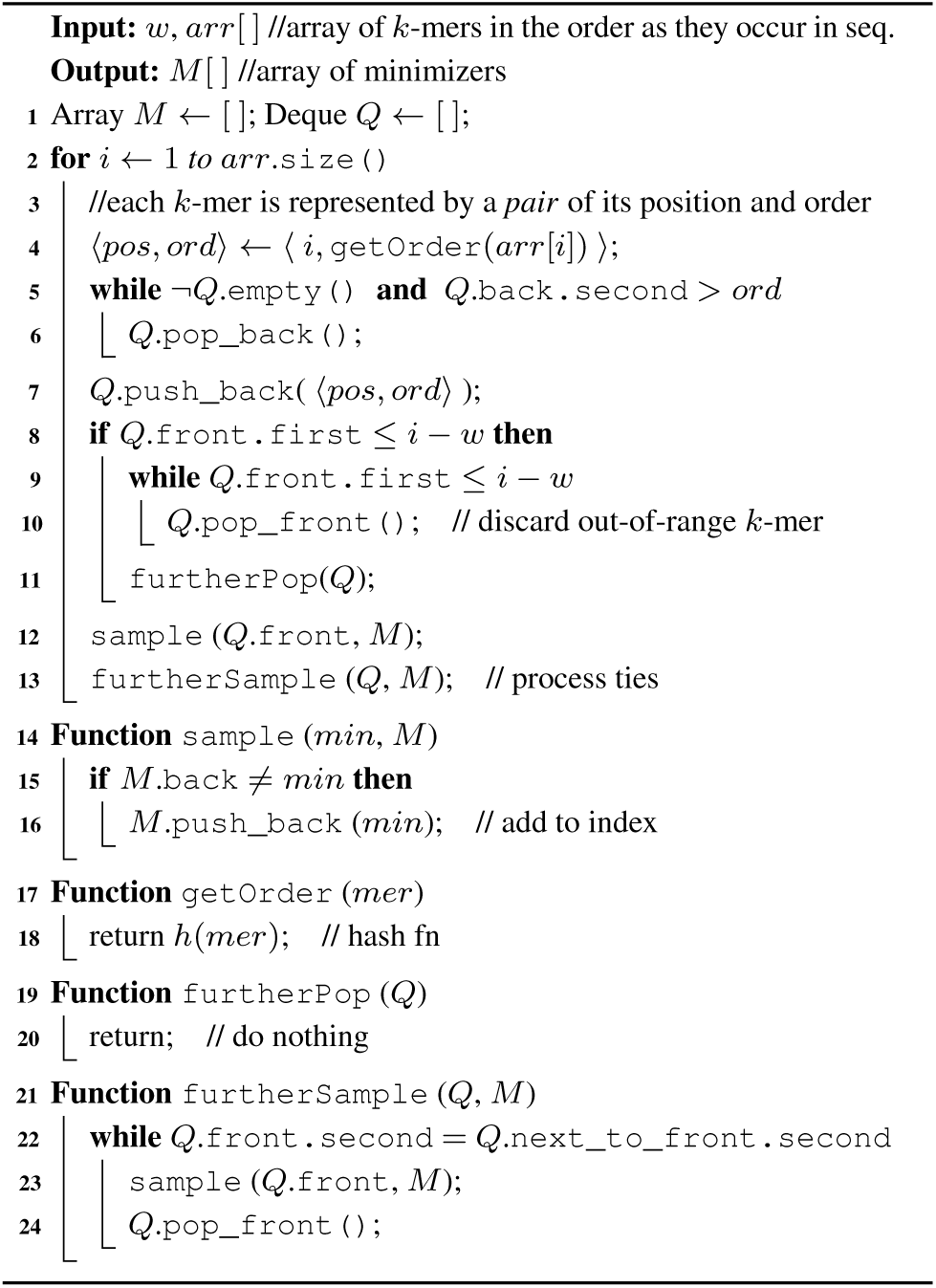

We intentionally split Algorithm 1 into a main routine and four helper functions. Later when we introduce optimizations to this algorithm to avoid sampling repetitive *k*-mers, we will only modify the functions leaving the main routine intact. Below, we summarize the properties of the minimizer sampling Algorithm 1 as a lemma. Readers are referred to the original articles (Roberts *et al.*, 2004; Smith, 2011) for formal proofs.

#### Lemma 1.

*The following statements are true:*

1. *Algorithm 1 computes the desired set of minimizers.*
2. *k-mer order values in queue Q of Algorithm 1 remain sorted in non-decreasing order during execution.*
3. *Assuming k-mer hashing is an O*(1) *operation, then Algorithm 1 uses O*(*n*) *time where n is length of input sequence.*
4. *If we ignore the possibility of k-mer ties within a window and assume that k-mer hash values are independent and uniformly distributed, then the expected density of minimizers is* 2*/*(*w* + 1).
5. *At least one matching minimizer will be sampled from two strings with matching substrings of length* ≥ *w* + *k* − 1.

### 2.2 MinHash and *k*-mer weighting

MinHash (Broder, 1997), like minimizer sampling, can be used to compute a signature of a sequence. It is a well-known locality sensitive hashing scheme for Jaccard similarity. Recall that the Jaccard similarity of two *k*-mer sets *X* and *Y* is |*X* ∩ *Y* |*/*|*X* ∪ *Y* |. Assuming *k*-mer hash values are independent and uniformly distributed, Broder (1997) proved that the probability of 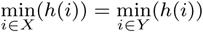 is equal to the Jaccard similarity. This enables an unbiased estimation of Jaccard while using only a subset of the original sets. When using Jaccard, elements in *X* ∩ *Y* contribute uniformly to the similarity value. However in some applications, the ‘importance’ of each element can vary. For instance, when comparing two sequences, a match of a rare *k*-mer carries more significance than a match of a highly-repetitive *k*-mer. Accordingly, Chum *et al.* (2008) proposed a variant of the Jaccard similarity metric called *weighted set-similarity*. Suppose function *µ* : *U* → ℕ assigns a weight to a *k*-mer, then the *weighted set-similarity* between two *k*-mer sets *X* and *Y* is given by:

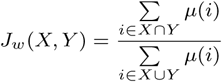

Note that *J*_*w*_ (*X, Y*) is more influenced by higher-weighted *k*-mers. It is possible to extend the MinHash algorithm to develop an unbiased estimator for *J*_*w*_. Let *h*_1_, *h*_2_ … *h*_*τ*_ be *τ* independent random hash functions where *τ* equals the highest *k*-mer weight 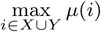. A *k*-mer *i* is hashed using the first *µ*(*i*) hash functions: 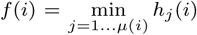. Naturally if a *k*-mer has a higher weight, its expected hash value *f* (*i*) is lower because more random hashes are considered. As a result, the probability of a *k*-mer *i* ∈ *X* being selected as signature of set *X* is decided by its weight and equals *µ*(*i*)*/*Σ_*j*∈*X*_ *µ*(*j*). Thus, similar to the original MinHash algorithm, the probability of 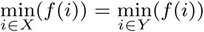 is *J*_*w*_ (*X, Y*).

In the above approach, we need to generate *µ*(*i*) uniformly distributed random numbers for each *k*-mer *i*. As a result, the above technique would require a large amount of space and time when input weights are large. Moreover, the above technique does not generalize to real-numbered weights. To remedy this, Chum *et al.* (2008) suggest an optimization which allows estimating *J*_*w*_ using a single random number for each *k*-mer. Their optimization follows from the observation that only the relative order of *k*-mer hash values matters and not their absolute values. As a result, it becomes possible to use a single random hash function while transforming its output to match the cumulative distribution of the hash function *f*. Their proposed alternative hashing function assigns *k*-mer *i* to 1 − (*h*_1_(*i*))^1*/µ*(*i*)^. This also generalizes the above algorithm to real-numbered weights *µ* : *U* → ℝ+. This algorithm partly inspired our formulation of weighted-minimizer sampling (Section 4).

## 3 Algorithm for Robust Winnowing

It is useful to recall the definition of *robust winnowing*:

### Definition 1

(Schleimer *et al.* (2003)). ***Robust winnowing:*** *In each window select the minimum k-mer. If possible break ties by selecting the same k-mer as the window one position to the left. If not, select the rightmost minimal k-mer.*

This tie breaking differs from the original algorithm of Roberts *et al.* (2004), which preserves all equally minimum *k*-mers. Robust sampling has one advantage and one disadvantage. The advantage is that we avoid sampling too many minimizers in low-complexity substrings such as “ACACAC…” or “AAAAA…” (Figure 3). Its disadvantage is that the minimizer selection from a window during a tie depends on the context, i.e. it depends on which minimizers were selected in prior windows. However, our results show that the advantage outweighs the disadvantage, and this optimization yields significant improvements in runtime and memory-usage without affecting accuracy.

**Fig. 3.**
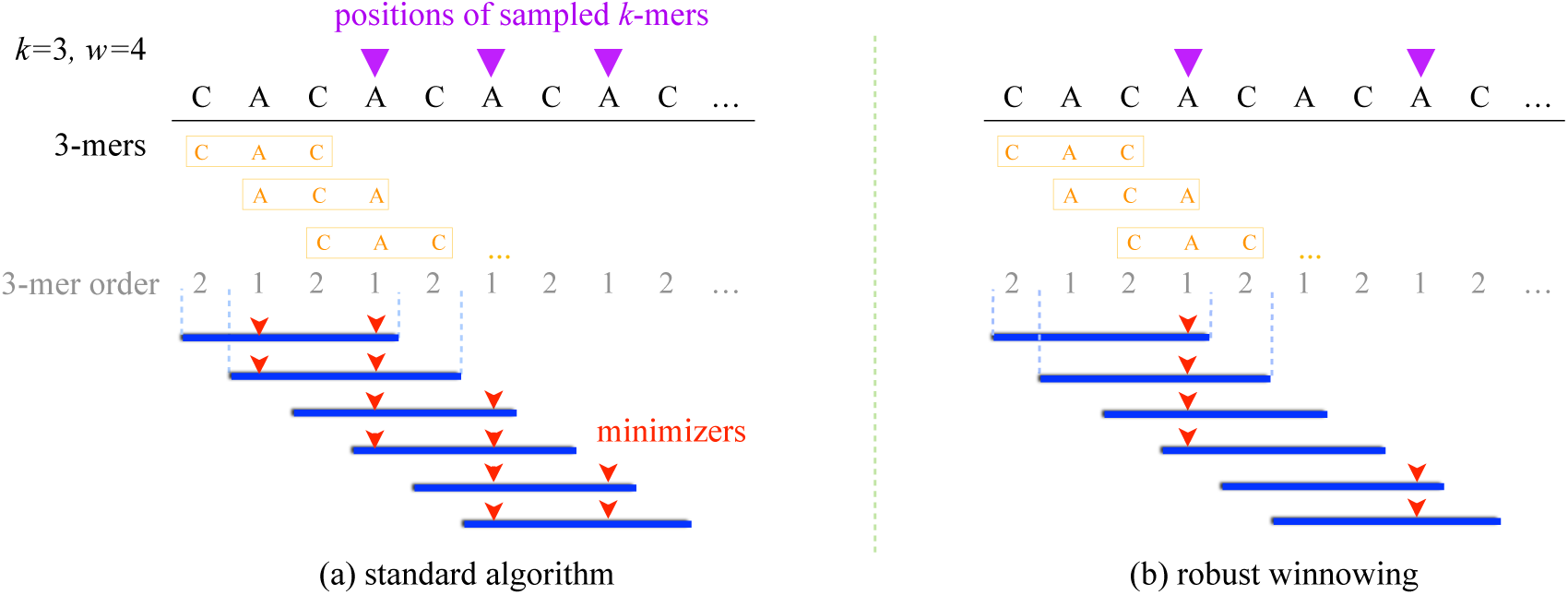
Visualization of tie breaking in the standard minimizer and robust winnowing algorithms. We use a low-complexity sequence “CACACA…” to illustrate the difference between the two approaches. The sequence is comprised of two 3-mers, “CAC” and “ACA”, which are ordered lexicographically in the above example. In the left plot, we break ties by sampling all equally-minimum *k*-mers per window, where as we follow the robust winnowing tie breaking in the right plot. Note that robust winnowing samples half the minimizers as compared to the standard approach in this repetitive sequence example.

Schleimer *et al.* (2003) did not provide an algorithm and time complexity for computing robust minimizers, so for completeness we provide here a linear-time algorithm (Algorithm 2) that borrows psuedocode from Algorithm 1, while modifying its `furtherPop` and `furtherSample` functions. Lemma 2 proves correctness of the proposed algorithm. Similar to Algorithm 1, it is trivial to argue that the runtime remains linear. Further, any matching substring of length ≥ *w* + *k* − 1 between two sequences sample at least one robust minimizer of same order.

### Algorithm 2: Computing minimizers using the robust winnowing method

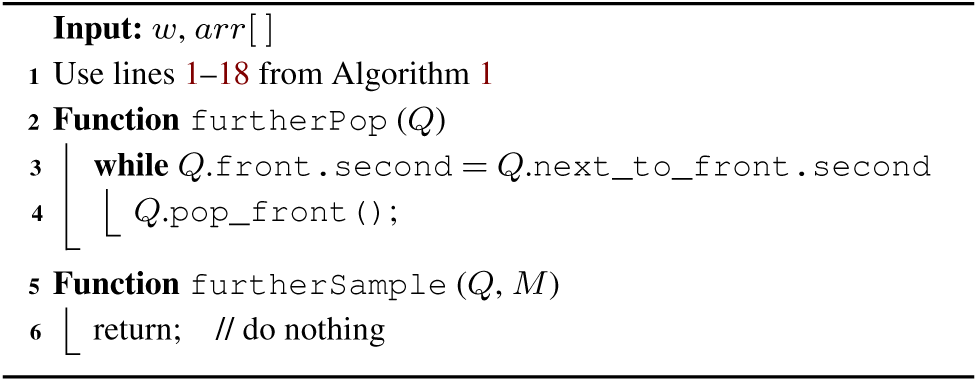

### Lemma 2. Algorithm 2 computes the desired set of minimizers using the robust winnowing criteria.

Proof. Using the robust winnowing scheme, we sample a new robust minimizer at iteration *i* iff either of the following are true: (i) the newest *k*-mer is lower than all *k*-mers in *Q*, or (ii) a prior robust minimizer (i.e. the front *k*-mer of *Q*) goes out of range. In the first case, the new *k*-mer will be selected as a robust minimizer by the `sample` function and pushed to the front of *Q*, while discarding other *k*-mers in *Q*. In the second case, we must pick the right-most minimum *k*-mer if there are multiple equal minimums in *Q*. The function `furtherPop` pops all but the right-most minimum in *Q*.

## 4 Weighted minimizer sampling

Using the same notation as before, function *µ* : *U* → ℝ+ assigns a weight to a *k*-mer. Weighted minimizer sampling is defined as following.

#### Definition 2.

***Weighted minimizer sampling:*** *Build a random ordering of all k-mers such that for a k-mer set X* = {*k*_1_, *k*_2_, …, *k*_|*X*|_} ⊆ *U, P* (min(*k*_1_, *k*_2_, …, *k*_|*X*|_) = *k*_*i*_) = *µ*(*k*_*i*_)*/*(*µ*(*k*_1_) + *µ*(*k*_2_) + … + *µ*(*k*_|*X*|_)) ∀*i* ∈ [1, |*X*|]. *Using this k-mer ordering, execute the robust-winnowing procedure.*

In the original minimizer sampling algorithm (Section 2.1), the *k*-mer ordering is defined directly using a random hash function. As a result, any *k*-mer in a window is equally likely to be selected as a minimizer (assuming no *k*-mer ties). The advantage of the weighted technique is that it allows us to bias selection of certain *k*-mers over others. If (*k*_1_, *k*_2_, … *k*_*w*_) is the set of *k*-mers in a window, then the probability of *k*-mer *k*_*i*_ being minimum equals *µ*(*k*_*i*_)*/*(*µ*(*k*_1_) + *µ*(*k*_2_) + … + *µ*(*k*_*w*_)). As a result, the higher the weight of a *k*-mer relative to its neighboring *k*-mers in a window, the more likely it is to be sampled as a minimizer. The weighted technique reduces to the original unweighted algorithm if all *k*-mer weights are equal. In order to define a *k*-mer ordering needed for weighted minimizer sampling, we borrow the optimized hashing technique of Chum *et al.* (2008) (Section 2.2). Given a random hash function *h* : *U* → [0, 1], we assign the order of a *k*-mer *k*_*i*_ to be 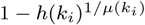.

We now analyze the expected density of weighted minimizer sampling. Recall that the expected density equals the expected fraction of *k*-mers sampled from a random input sequence. While the original algorithm guarantees density of 2*/*(*w* + 1) (Lemma 1), here the density depends on *k*-mer weights.

### Lemma 3.

*Suppose k*_1_, *k*_2_, … *k*_*n*−*k*+1_ *are the k-mers in the order as they appear in a random input sequence of length n. Assume σ*^*k*^ ≫ *w. The expected density of weighted minimizer sampling equals:*

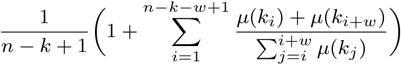

Proof. Suppose *W*_*i*_ is window of *w* consecutive *k*-mers starting at position *i*. Window *W*_*i*_ is said to be *charged* if the minimum *k*-mer in window *W*_*i*_ differs from the minimum *k*-mer in window *W*_*i*−1_, i.e. *W*_*i*_ selects a new minimum. The expected density of a minimizer sampling can be analyzed by counting total number of charged windows in the input sequence.

A union of two windows *W*_*i*_ and *W*_*i*−1_ defines an interval of length *w* + 1 in the input sequence. Accordingly, *k*-mers *k*_*i*−1_, *k*_*i*_, … *k*_*i*+*w*−1_ occur in this interval, and let *k*_*min*_ be the minimum *k*-mer among these. As *σ*^*k*^ ≫ *w*, we safely ignore the possibility of ties. Window *W*_*i*_ is charged if and only if either *k*_*min*_ = *k*_*i*−1_ or *k*_*min*_ = *k*_*i*+*w*−1_. Define random variable *X*_*i*_ to be 1 if *W*_*i*_ is charged and 0 if not.

For *i* ∈ [2, *n* − *k* − *w* + 2] (for all windows except the first)

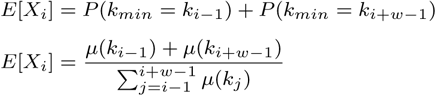

For *i* = 1, *E*[*X*_*i*_] = 1. Therefore, the expected density

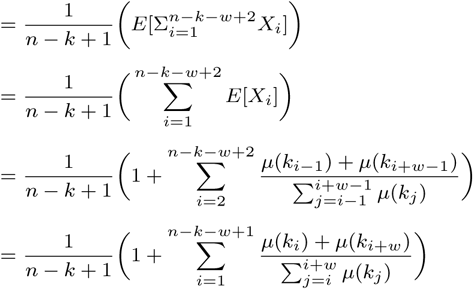

The theoretical worst case expected density of weighted minimizer sampling occurs when *k*-mers *k*_1_, *k*_2_, …, *k*_*n*−*k*+1_ are distinct (i.e., *σ*^*k*^ ≫ *n*) and *µ*(*k*_1_) ≫ *µ*(*k*_2_) ≫ … ≫ *µ*(*k*_*n*−*k*+1_). In this case, the density approximately equals 1. A best case scenario is one where *k*_1_, *k*_2_, …, *k*_*n*−*k*+1_ are distinct and weights *µ*(*k*_*w*_), *µ*(*k*_2*w*_), *µ*(*k*_3*w*_) … are significantly higher than all the other *k*-mer weights. In this case, density is optimal and equals 1*/w*.

Winnowmap implements weighted minimizers sampling to avoid indexing highly repetitive *k*-mers for long-read mapping. Given any reference genome, we assume the availability of a pre-computed set 𝒮 of repetitive *k*-mers, i.e. *k*-mers that occur with frequency above a certain cutoff. This set can be computed efficiently using a *k*-mer counting tool. We assign any *k*-mer *k*_*i*_ ∈ *U* a weight of *v <* 1 if *k*_*i*_ ∈ 𝒮, and 1 otherwise (Algorithm 3). By default, Winnowmap uses a frequency cutoff of 1024 for determining highly repetitive *k*-mers, and sets *v* = 1*/*8. Note that the asymptotic time complexity of minimizer sampling remains linear in sequence length if we use a *σ*^*k*^-sized bit array to store the set 𝒮 that supports membership queries in *O*(1) time.

### Lemma 4.

*The following statements are true for Algorithm 3:*

1. *Assuming k-mer hashing is an O*(1) *operation, then Algorithm 3 uses O*(*n*) *time where n is length of input sequence.*
2. *At least one minimizer of the same order will be sampled from two strings with matching substrings of length* ≥ *w* + *k* − 1.

Algorithm 3 involves computational overhead that affects indexing performance in practice. First, random accesses to a *σ*^*k*^-sized bit array in each iteration makes the implementation slow (line 5). Second, computing a power arithmetic operation for each occurrence of repetitive *k*-mers is expensive (line 6). We implement two optimizations to avoid this overhead. As |𝒮| is expected to be ≪ *σ*^*k*^, we use a space-efficient bloom filter to query membership in 𝒮, which incurs a marginal false-positive rate. For the second issue, we compute *x*^1*/v*^ without using a power operation. For example, if *v* is 1*/*8, then *x*^1*/v*^ = *x*^8^ can be computed using three multiplications. Finally, we use −*x*^1*/v*^ (line 6) and −*x* (line 7) to assign *k*-mer order values instead of 1 − *x*^1*/v*^ and 1 − *x* respectively to avoid rounding errors (e.g., 1 − 1e-20 rounded to 1). As a result of these optimizations, weighted minimizer sampling is nearly as fast as the standard minimizer sampling.

### Algorithm 3: Algorithm for computing weighted minimizers

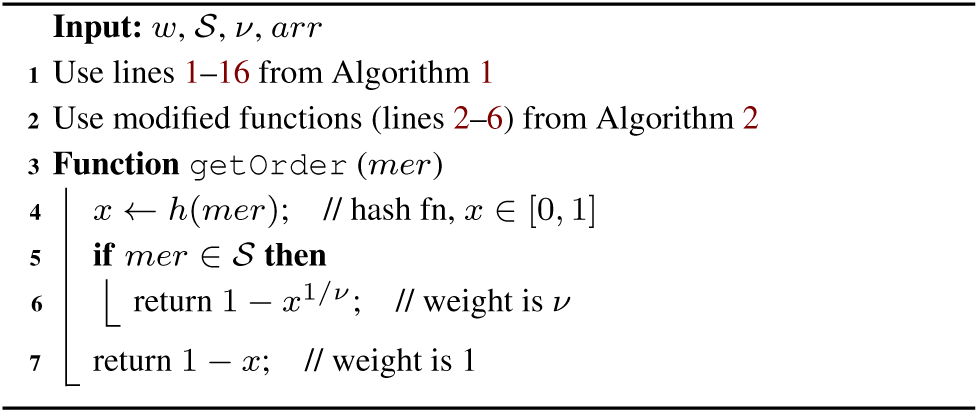

## 5 Results

In this section, we demonstrate the empirical advantage of robust winnowing and weighted minimizer sampling for long read mapping. As mentioned earlier, we have implemented Winnowmap by replacing the minimizer sampling and indexing procedures of the well-established mapping tool Minimap2 (Li, 2018). Minimap2 has been previously shown to achieve excellent read mapping results in terms of accuracy and speed, so in this section we compare Winnowmap directly against Minimap2 (v2.17-r954-dirty).

### 5.1 Experimental setup

#### Hardware and software

For all our experiments, we used a server equipped with two Intel Xeon E5-2698 v4 20-core processors and 1 TB memory. Both Winnowmap and Minimap2 were run in parallel-mode using 16 CPU threads, and we report wall-clock time in our results. We used recommended parameters for mapping PacBio and Oxford Nanopore (ONT) reads based on Minimap2’s user documentation. These are `-cx map-pb` for PacBio reads, and `-cx map-ont` for ONT reads. Winnowmap requires an additional parameter specifying a file containing the list of highly-repetitive *k*-mers for a reference genome. In our experiments, this file was assumed to be pre-computed. Given the latest advances in *k*-mer counting algorithms, this is a low-overhead *O*(*n*) operation, especially for the values of *k* typically used for long-read mapping (e.g. 15 for ONT reads and 19 for PacBio reads).

Winnowmap uses a default window length of 50. In contrast, Minimap2 performs dense minimizer sampling using a default window length of 10. Our results show that, unlike Minimap2, Winnowmap can tolerate sparser sampling without compromising accuracy. Minimap2 relies on standard minimizer sampling (Roberts *et al.*, 2004) before masking the top 0.02% most frequent minimizers from its index. Winnowmap does not use any masking heuristics. All other algorithmic features and parameters are the same between Minimap2 and Winnowmap.

#### Benchmarking data sets

Our data sets include two simulated sets of PacBio reads (D1, D2) and one set of real ultra-long ONT reads (D3). Total count, N50, and maximum length of these read sets is shown in Table 5.1. The simulated read sets were obtained using PBSIM (Ono *et al.*, 2013) with a mean error-rate of 10% and mean read length of 15 kbp. PacBio read set D1 was simulated from a recently finished human X chromosome (Miga *et al.*, 2019) and D2 simulated from the human whole-genome reference GRCh38 (Schneider *et al.*, 2017), respectively. The assembly of chromosome X (v0.7) contains the first-ever resolved centromere (3.1 Mbp), which allowed us to evaluate mapping accuracy within a long tandem repeat. Finally, read set D3 is a small random sample of the `rel3` human ‘CHM13’ whole-genome, ultra-long ONT sequencing data (Miga *et al.*, 2019). A minimum read length of 1 kbp was enforced during sub-sampling. The read sets were mapped to the human chromosome X assembly (D1) and the human reference genome (D2, D3), respectively.

#### Evaluation criteria

For simulated read sets D1 and D2, we evaluated mapping accuracy against the known truth. We followed a similar criteria previously used by Li (2018). A read is said to be mapped correctly if its primary alignment overlaps with the true interval, and the overlapping bases constitute ≥ 10% of the union of the true and the reported alignment intervals. We used Minimap2’s `paftools` utility (Li, 2018) to compute mapping accuracy using this criteria. We also measured indexing time, mapping time, peak memory-use, fraction of unmapped reads, and minimizer sampling density.

### 5.2 Comparison with Minimap2

We evaluated performance of Winnowmap and Minimap2 using data sets D1–D3 (Table 5.1). We tested the two algorithms using both window length parameters 10 and 50. While discussing differences between Winnowmap and Minimap2, we refer to the results obtained using their default window length parameters, i.e. 50 for Winnowmap, 10 for Minimap2. Using simulated read sets D1 and D2, Winnowmap achieves better read mapping accuracy than Minimap2. Moreover, the accuracy of Winnowmap is largely unaffected by the varying window length. Compared to Minimap2, Winnowmap reduces read mapping error from 0.14% to 0.06% on data set D1, and from 1.85% to 1.81% on data set D2. Both Winnowmap and Minimap2 were able to map all reads in D1 and D2. In the D3 data set, there are many very long reads but also a higher fraction of short reads. This resulted in a significant fraction of unmapped reads using both algorithms. Using its default window length, Winnowmap has comparable mapping and indexing runtime to Minimap2, while using less memory. However, with a small window length of 10, the runtime and memory use of Winnowmap is much higher because weighted minimizer sampling is less effective for small window lengths. Large windows provide more freedom for the weighted minimizer sampling to choose a non-repetitive *k*-mer over a repetitive *k*-mer.

Winnowmap significantly improves accuracy in large repeats. Using data set D1, we evaluated its accuracy within the centromeric region of chromosome X, which comprises approximately 1500 tandemly arrayed copies of a 2057 bp repeat unit spanning approximately 3.1 Mbp. To measure mapping accuracy within this repeat array, we considered all 1057 reads that were sampled from the region (positions 57, 828, 561 to 60, 934, 693). Table 3 shows the mapping accuracy with different window length parameters. Compared to Minimap2, Winnowmap reduces mapping error from 3.6% to 0.0% and its accuracy is again robust to changing window length. This improvement is possible since Winnowmap maintains all minimizers in the index whereas Minimap2 masks repetitive minimizers. To visualize the effect of masking, we show minimizer sampling density across the length of chromosome X in Figure 4. This plot shows the minimizer sampling density of three methods: (i) the standard minimizer sampling (Roberts *et al.*, 2004) without any modification (blue), (ii) minimizer sampling and masking in Minimap2 (orange), and (iii) weighted minimizer sampling in Winnowmap (green). The window length was fixed to 10 for all the three methods. Minimap2’s masking heuristic leads to poor sampling density in long repeats with the most significant dip observed in the centromere region.

**Table 1.**
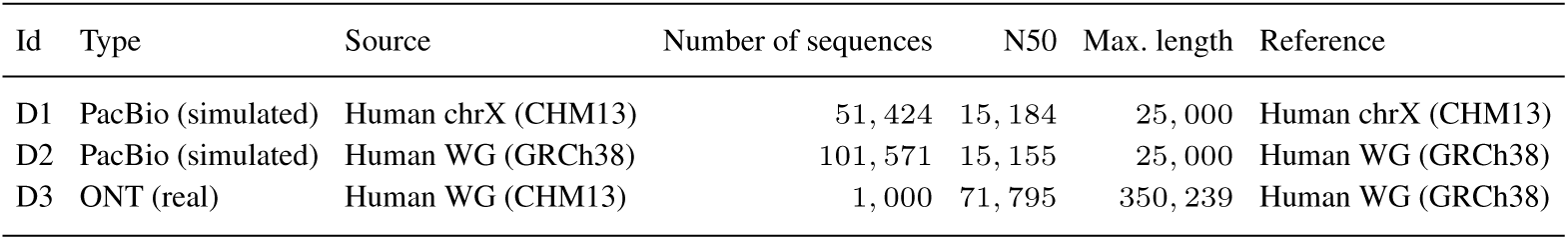
List of datasets used for evaluation.

**Table 2.** Comparison of mapping performance between Winnowmap and Minimap2 using data sets D1–D3. We tested both methods while varying window length, which controls the minimizer sampling rate. Larger window lengths lead to more sparse sampling, as indicated by the last column. Minimap2 uses *w* = 10 by default. Winnowmap is more robust to sparse sampling and, as a result, uses a larger default window length of 50.

| Data set | Window length ($w$) | Method | Unmapped reads | Incorrectly mapped reads | Indexing time (s) | Mapping time (s) | Memory-usage (GB) | Average minimizer spacing |
| --- | --- | --- | --- | --- | --- | --- | --- | --- |
| D1 | 10 | Winnowmap | 0.00% | 0.06% | 5.5 | 107.6 | 8.5 | 7.8 |
|  | 50 | Winnowmap | 0.00% | 0.06% | 6.7 | 34.8 | 2.9 | 36.2 |
|  | 10 | Minimap2 | 0.00% | 0.14% | 4.9 | 26.5 | 5.9 | 7.8 |
|  | 50 | Minimap2 | 0.01% | 0.77% | 3.0 | 32.8 | 5.9 | 36.3 |
| D2 | 10 | Winnowmap | 0.00% | 1.83% | 67.6 | 1292.6 | 91.2 | 8.2 |
|  | 50 | Winnowmap | 0.00% | 1.81% | 57.7 | 92.2 | 7.3 | 37.6 |
|  | 10 | Minimap2 | 0.00% | 1.85% | 55.4 | 70.3 | 11.1 | 8.2 |
|  | 50 | Minimap2 | 0.02% | 2.07% | 40.1 | 67.7 | 8.7 | 38.2 |
| D3 | 10 | Winnowmap | 4.50% | NA | 87.4 | 2901.0 | 70.4 | 5.7 |
|  | 50 | Winnowmap | 10.80% | NA | 65.1 | 39.8 | 9.9 | 26.3 |
|  | 10 | Minimap2 | 9.70% | NA | 62.6 | 6.7 | 12.5 | 5.6 |
|  | 50 | Minimap2 | 14.70% | NA | 40.2 | 3.1 | 7.3 | 25.7 |

**Table 3.** Evaluation of mapping accuracy in the centromeric region of human chromosome X. Here we consider only those reads in data set D1 which were sampled from centromeric repeat array.

| Window length ($w$) | Method | Unmapped reads | Incorrectly mapped reads |
| --- | --- | --- | --- |
| 10 | Winnowmap | 0.0% | 0.0% |
|  | Minimap2 | 0.0% | 3.6% |
| 25 | Winnowmap | 0.0% | 0.0% |
|  | Minimap2 | 0.0% | 14.5% |
| 50 | Winnowmap | 0.0% | 0.0% |
|  | Minimap2 | 0.6% | 34.7% |

**Fig. 4.**
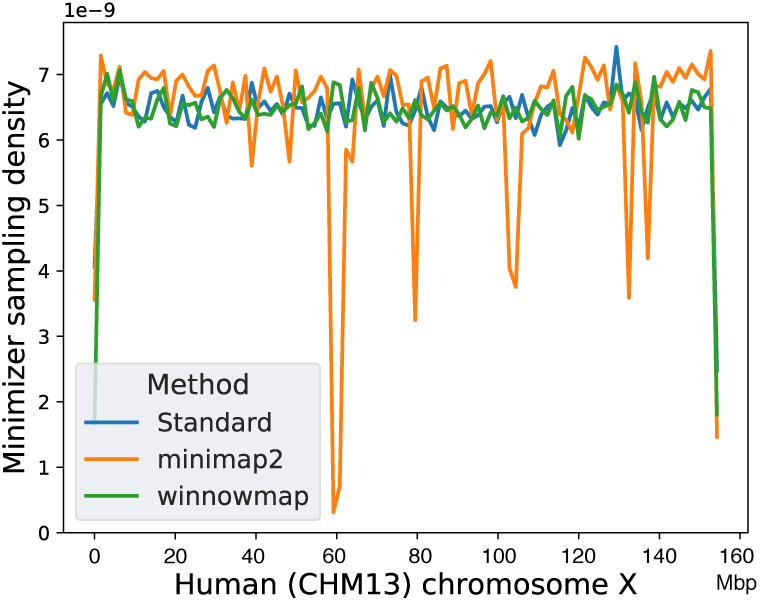
Minimizer sampling density using a complete human chromosome X as the reference (data set D1). We compare three sampling algorithms using fixed window length *w* = 10. ‘Standard’ method refers to the classic minimizer sampling algorithm from Roberts et al. (2004), without any masking or modification. Minimap2 uses the standard algorithm, but masks the most frequently occurring minimizers (top 0.02%) in the reference (count ≥ 160 for this reference). Winnowmap uses weighted minimizer sampling. Both ‘Standard’ and Winnowmap methods maintain at least one minimizer per window in their index and achieve near-uniform density. The masking heuristic in Minimap2 affects long repetitive segments (e.g. centromere: 58–61 Mbp), where density drops significantly.

### 5.3 Benefit of weighted minimizer sampling and robust winnowing

Weighted minimizer sampling in Winnowmap includes two key optimizations, (i) robust winnowing, and (ii) weighted sampling. These two optimizations are independent of one another and we evaluated the advantages of each. To do this, we compared (i) Winnowmap, (ii) a version of Winnowmap that implements only robust winnowing, and (iii) a version of Minimap2 that implements the standard minimizer sampling without the masking heuristic. Accordingly, these three methods are referred to as ‘Optimization 1+2’, ‘Optimization 1’, and ‘Standard’, respectively in Figure 5. As there is no masking involved, these three methods index all minimizers irrespective of their frequency. Here we show mapping time and peak memory usage using data sets D1–D3 while keeping window length equal. We used a window length of 10 for data sets D1 and D2, and 50 for data set D3. The ‘Standard’ method crashed on data set D3 with a window length of 10. These results demonstrate that these Winnowmap optimizations are crucial for speed and memory efficiency.

**Fig. 5.**
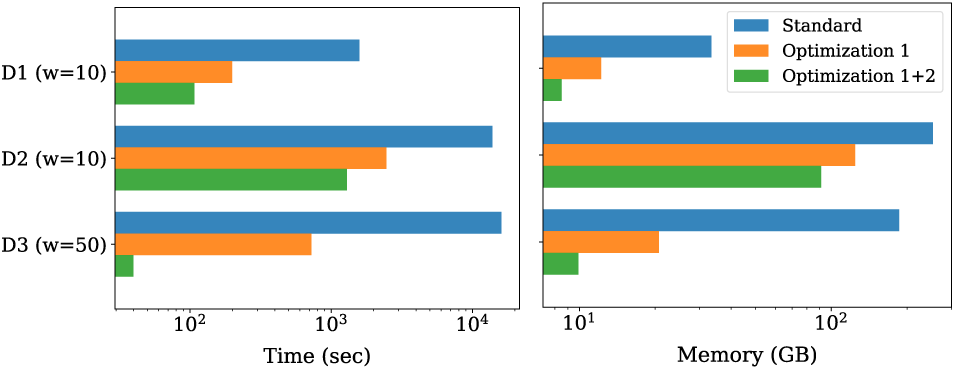
Benefit of the proposed minimizer sampling optimizations to long read mapping performance. We compare three methods: ‘Standard’ (default minimizer sampling, no masking), ‘Optimization 1’ (robust winnowing) and ‘Optimization 1+2’ (robust winnowing and weighted minimizer sampling). All three methods select at least one minimizer in each window. The left plot indicates total mapping time whereas the right plot indicates total memory usage, using data sets D1–D3. The x-axis is log-scaled in both plots. The combination of both optimizations is crucial in Winnowmap to achieve good efficiency.

## 6 Conclusions

Minimizer sampling is a simple yet powerful technique to speed up genome sequence analysis. When this technique is used for seeding sequence alignments, minimizers with high frequency often lead to too many false-positive seed matches. The large number of false hits naturally leads to high memory-usage and runtime. To date, a popular way to deal with this issue has been to mask highly repetitive minimizers. While this masking heuristic eliminates false hits, the underlying guarantee of a minimizer seed match is lost in the process. As a result, masking decreases mapping accuracy within long genomic repeats. In this paper, we describe optimizations to the standard minimizer sampling procedure and evaluate the improvements to long-read mapping. We have introduced weighted minimizer sampling, where users can specify which *k*-mers should be more (or less) likely to be selected as minimizers. Lastly, when there are multiple equally minimum *k*-mers in a window (e.g. in low complexity regions), robust winnowing is effective in preventing oversampling.

We implemented weighted minimizer sampling and robust winnowing in Winnowmap. Winnowmap makes it feasible to map PacBio or ONT reads without the need for a masking heuristic. As a result, it achieves superior mapping accuracy by maintaining a uniform sampling density across the reference sequence using a simple weighting criteria.

Future work will be focused in three directions. First, we will explore more sophisticated weight functions, e.g. down-weighting erroneous *k*-mers in addition to repetitive *k*-mers. A set of likely erroneous *k*-mers could be detected prior to mapping by intersecting the *k*-mer sets of the reads and the reference genome. Second, we will test the idea of weighted minimizer sampling in other applications such as long-read overlapping and metagenomic read classification. Third, the density bounds that were discussed in this paper are defined for arbitrary weights. However, in most applications, we can assume weight values are bounded between [1*/c, c*], where *c* is a constant. It will be useful to derive tight bounds on the expected density under this constraint.

